# Microbial diversity and toxin risk in tropical freshwater reservoirs of Cape Verde

**DOI:** 10.1101/266957

**Authors:** Ana P. Semedo-Aguiar, José B. Pereira-Leal, Ricardo B. Leite

## Abstract

The Cape Verde islands are part of the African Sahelian arid belt that possesses an irregular rainy season between August and October. This erratic rain pattern has prompted the need for water reservoirs, now critical for the country’s sustainability. Worldwide, freshwater cyanobacterial blooms are increasing in frequency due to global climate change and eutrophication of water bodies, particularly in reservoirs. To date there have been no risk assessments of cyanobacterial toxin production in these man-made structures. We evaluated this potential risk using 16S rRNA gene amplicon sequencing and full metagenome sequencing in freshwater reservoirs of Cape Verde.

Our analysis revealed the presence of several potentially toxic cyanobacterial genera in all sampled reservoirs (Poilão, Saquinho and Faveta). In Faveta *Microcystis* sp., a genus well known for toxin production and bloom-formation, dominated our samples, while a green algae of the genus *Cryptomonas* and *Gammaproteobacteria* dominated Saquinho and Poilão.

Taking advantage of the dominance of *Microcystis* in the Faveta reservoir, we were able to reconstruct and assemble its genome, extracted from a metagenome of bulk DNA from Faveta water. We named it *Microcystis* cf. *aeruginosa* CV01, for which a phylogenetic analysis revealed to have a close relationship with other genomes from those taxa, as well as other continental African strains, suggesting geographical coherency. In addition, it revealed several clusters of known toxin-producing genes. This assessment of Cape Verdean freshwater microbial diversity and potential for toxin production reinforces the need to better understand the microbial ecology as a whole of water reservoirs on the rise.

## Introduction

The available freshwater in the African archipelago of Cape Verde (DMS coordinates 15°07′12.51” N, 23°36′18.62” W) does not cover its needs. In addition, overexploitation and saline intrusion can impair the quality of groundwater [1]. Several water storage structures are being planned around the country in order to collect storm water needed to increase irrigated areas and modernize agriculture. However, these waters can carry nutrients of natural and anthropogenic origin, creating conditions for eutrophication and exponential growth of microalgae. These algal blooms have deleterious impacts on public health, water quality, and environmental issues, as well as economic costs due to bottom anoxia, release of noxious products, and toxic metabolites [2,3]. Actually, these events are occurring more frequently worldwide, and it is thought that global climatic changes are a major contributor to this problem [4–6].

To evaluate the risk of occurrence of algal blooms in freshwater bodies it is important to characterize their microbial composition. In non-eutrophic freshwater systems, the most commonly abundant bacterial groups are, in order of decreasing relative abundance, *Actinobacteria*, *Bacteroidetes*, *Proteobacteria* (Alpha, Beta and Gamma clades), *Verrucomicrobia* and *Cyanobacteria* [7–12]. In contrast, eutrophic waters contain microbial communities that include large numbers of cyanobacteria, which are able to produce toxins and foul odors and discolor the water [13,14]. The cyanobacterial phylum has many genera that produce toxins, also called cyanotoxins, and in freshwater bodies, toxic and non-toxic strains can co-exist and dominate at different times [15,16]. Poisoning with cyanotoxins occurs through consumption of contaminated food or water, or during aquatic recreational activities, causing many adverse symptoms like skin irritation, acute diarrhea, and liver and nervous tissue damage, leading to severe health problems, or death in humans, cattle, domestic animals and wildlife [6,13,17–19]. Hence, the risk of having toxin production increases the need for monitoring plans to prevent toxin-related impairments and costs.

Molecular-based methods combined with sequencing offer the ability to not only identify possible toxin producers but also target species-specific toxins, validating the presence or absence of toxin-related pathways. Nevertheless, DNA-based molecular methods cannot predict if toxins are being produced and released to the environment.

The leveraging of molecular methods provided by Next Generation Sequencing (NGS) allows researchers to gain new insights into microbial community structure in environment samples, identify new community members and discover potential bioindicators of water quality.

To determine the current risk of toxin production in the Cape Verdean freshwater reservoirs, we performed NGS analysis of 16S rRNA gene amplicon sequences to identify microalgae and bacteria in the reservoirs. Our results show the presence of several cyanobacterial genera well known to produce toxins in all reservoirs. We were able to reconstruct, for the first time, the full genome of a potentially toxic cyanobacterium from Cape Verde, based on the full metagenome sequencing data of Faveta reservoir. Analysis of this genome revealed the presence of genetic machinery used to synthesize cyanotoxins. The results of our biodiversity survey, phylogenetic analysis, and genome reconstruction, lead us to conclude that toxin risk is a reality and potential future threat in these reservoirs.

## Materials and methods

### Study sites and Sampling

This study was conducted on the Cape Verdean Island of Santiago (Fig 1). During sampling, in February 2014, only Poilão (B1), Saquinho (B2) and Faveta (B3) reservoirs had water while Figueira Gorda and Salineiro were empty at the tie of sampling. Other contextual information on Santiago Island’s reservoirs is summarized in Table 1.

**Fig 1.**
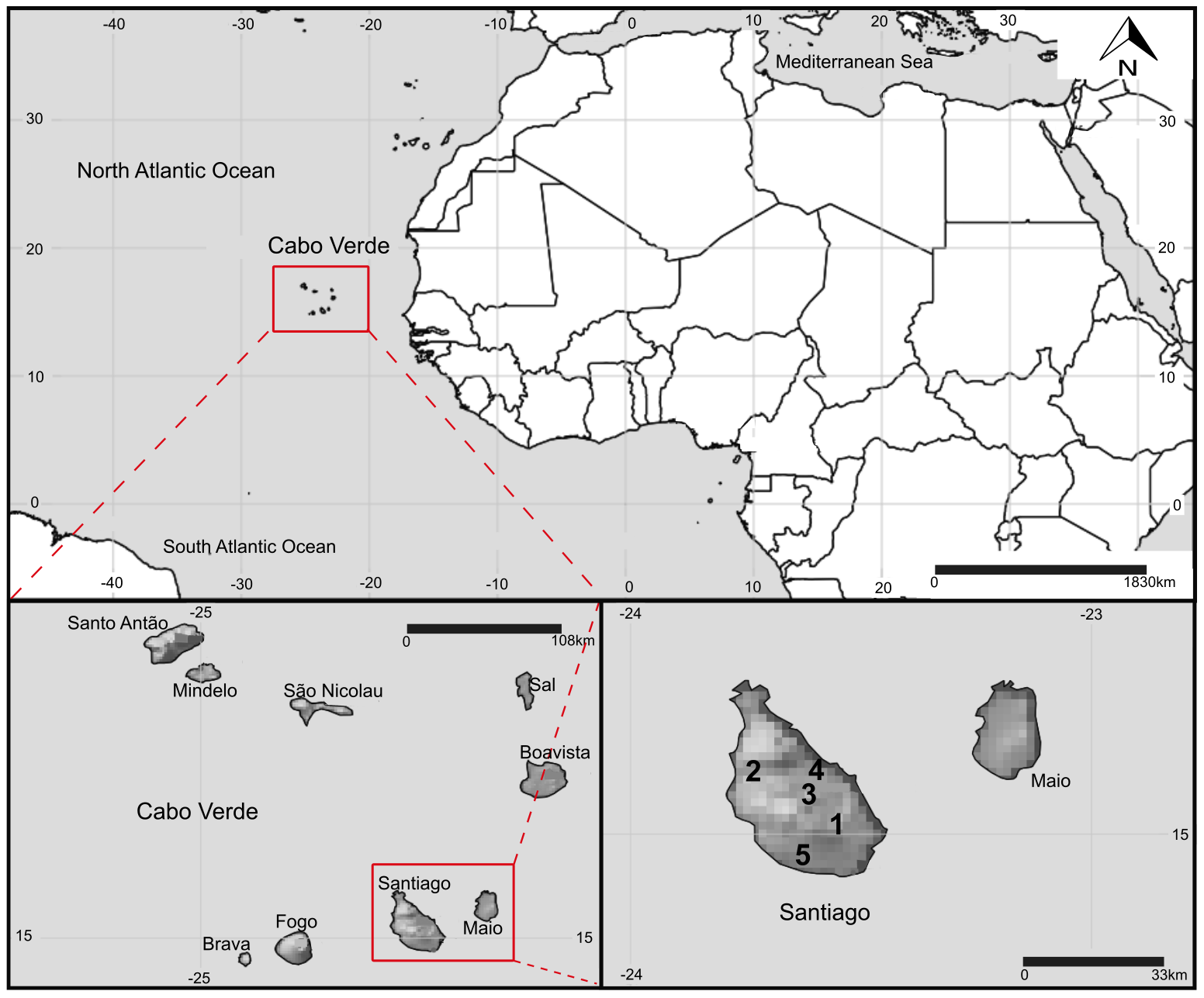
Map of Cape Verde and location of the reservoirs in Santiago Island sampled in this study. DMS coordinates of the reservoirs: 1-Poilão (15°04’25.0”N, 23°33’25.8”W), 2-Saquinho (15°08’11.3”N, 23°42’27.7”W) and 3-Faveta (15°05’54.5”N, 23°37’24.1”W). 4-Figueira Gorda (15°07’5.8”N, 23°35’36.5”W) and 5-Salineiro (14°57’03.5”N, 23°38’00.4”W) reservoirs where empty at the time of sampling.

**Table 1.**
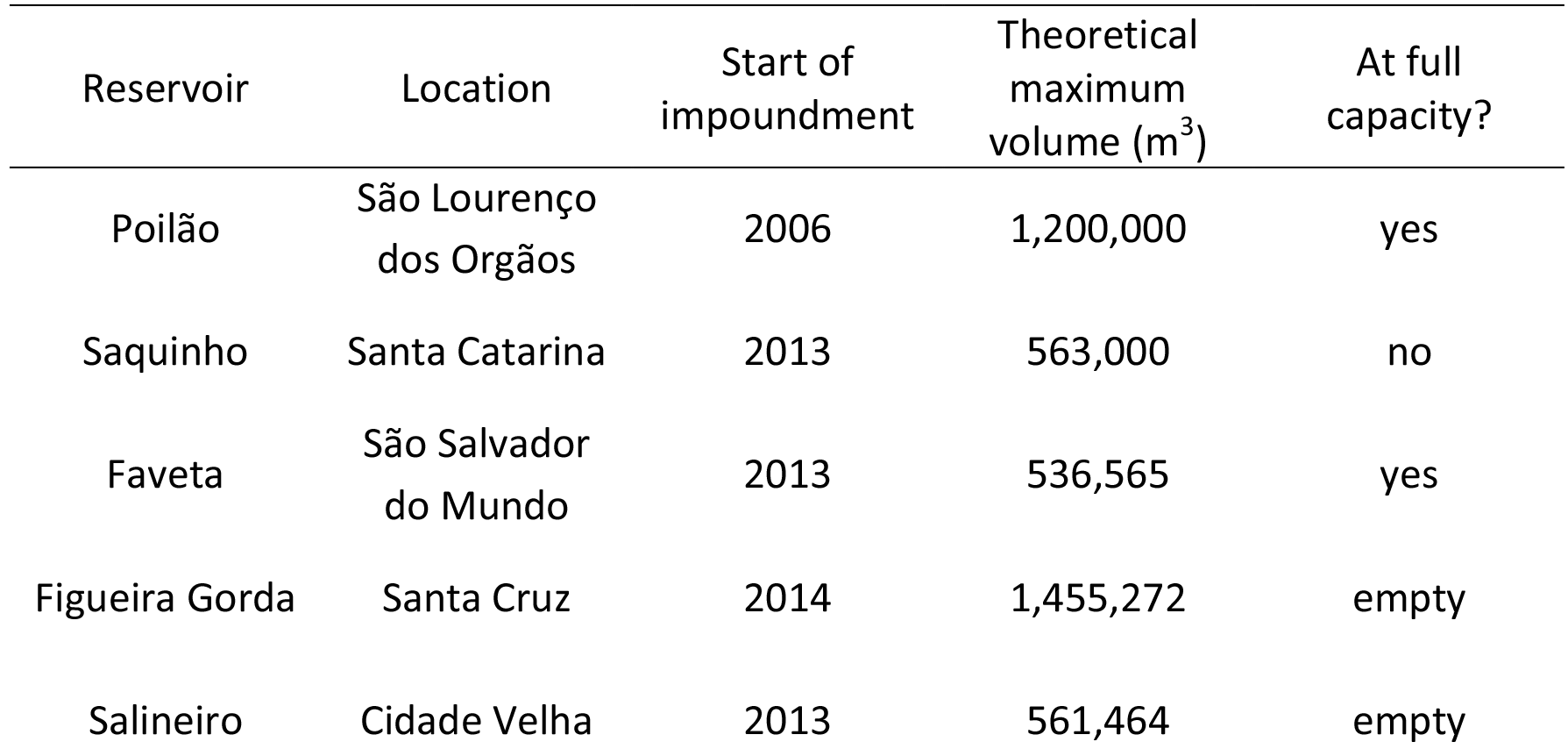
Characteristics of the studied reservoirs in February 2014 (Government of Cape Verde)

| Reservoir | Location | Start of impoundment | Theoretical maximum volume (m<sup>3</sup>) | At full capacity? |
| --- | --- | --- | --- | --- |
| Poilão | São Lourenço dos Orgãos | 2006 | 1,200,000 | yes |
| Saquinho | Santa Catarina | 2013 | 563,000 | no |
| Faveta | São Salvador do Mundo | 2013 | 536,565 | yes |
| Figueira Gorda | Santa Cruz | 2014 | 1,455,272 | empty |
| Salineiro | Cidade Velha | 2013 | 561,464 | empty |

Triplicate water samples were collected from 0.5 to 1 m depth at each study site in the same sampling area, using a five-liter bucket. Microbial cells were concentrated by filtering the five liters of water through 0.22 μL Sterivex-GV filters (Millipore, Billerica, MA, USA) by vacuum filtration. Filters where kept on ice and in the dark while transported to the lab for DNA extraction.

### DNA extraction

Genomic DNA was extracted from the individual filters using PowerWater Sterivex DNA Isolation Kit (MO BIO, USA) following the manufacturer’s protocol. The amount of the DNA extracted was later quantified using NanoDrop 1000 spectrophotometer (Thermo-Fisher Scientific, Wilmington, DE, USA) measuring the UV absorption at 260 nm and 280 nm wavelengths.

### Bacterial diversity: 16S rRNA gene amplicon sequencing

To assess the bacterial diversity in the Cape Verdean reservoirs, we used the 16S rRNA gene as a marker for biodiversity.

The extracted environmental DNA was amplified using primers targeting the V3 and V4 hypervariable regions of the 16S rRNA gene. PCR amplification of the 464 bp fragments was performed with the general bacterial primer pair 341F/785R [20].

The purified DNA was sequenced on an Illumina MiSeq platform using a 250 bp paired-end DNA library to generate at least 100,000 reads per sample. 16S ribosomal RNA gene amplicon sequencing was performed by Instituto Gulbenkian de Ciência (IGC) Gene Expression Facility.

### Taxonomic composition and identification of potentially toxic genus/species

We used the QIIME v1.9.1 workflow [21] for demultiplexing, removing barcodes, quality filtering, clustering, and taxonomy assignment of the reads. Briefly, merging of the paired-end reads was done with a defined minimum overlap of 8 bp and a maximum difference of 20%, followed by quality filtering of the reads using default parameters [22], for each reservoir. Sequences were then clustered at a 97% identity threshold and all clusters with less than 4 sequences were removed thus reducing noisy reads. For each cluster a reference sequence was chosen (the first) and by comparison with Greengenes v13_8 [23], using UCLUST algorithm [24] with default parameters into Operational Taxonomic Units - OTUs. Taxonomy assignments were done with RDP Classifier v2.2 [25]. To assess the consistency of the sampling and the biodiversity present in each sampling site, diversity indices where calculated: number of Observed species, Shannon’s index and Simpson’s Dominance index [26].

### Graphical representation of the microbial communities’ structure detected in the reservoirs

Phyloseq package [27] in RStudio 1.1.419 [28] was used to perform a graphical representation of the structure of the microbial community in each of the sampled sites. A complete assignment of the more abundant bacterial genera in our samples, was generated by Kraken running the full Bacterial Genbank Reference sequences [29], excluding the sequences assigned previously as chloroplasts by QIIME.

### Sequencing and assembly of the *Microcystis* cf. *aeruginosa* CV01 genome

A whole metagenome sequencing of Faveta reservoir was performed using Illumina MiSeq v3 kit with a 200x coverage. The DNA libraries were created using Nextera DNA Library Preparation Kit, and sequencing was executed by IGC’s Gene Expression Facility.

The sequenced reads were assembled with SPAdes v3.6.0 [30] using single-cell mode, and k-mers of 21, 33, 55, 77, 99 and 127. The GC content of the assembled metagenome from Faveta reservoir revealed two distinct contributions: one clearly in the range of the GC content typical of *Microcystis aeruginosa* (42-43%) [31] and the other peak, at approximately 63% for other contributors present in the sample, such as several GC rich strains of *Gamma*, *Alpha,* and *Betaproteobacteria* (S1 Fig). The peak with lower GC was in the range of our target species and most of its reads have high coverage numbers since they belong to the numerically most abundant species. Two different approaches were used to isolate this genomic contribution: one using CONCOCT v0.41 [32] a binning tool that operates based on sequence coverage and composition, and another exploring the high coverage percentage of a single genomic contribution by assembling reads from the raw sequencing data that had coverage above 60x. In both methods, short sequences were filtered out (length < 1,000 bp). Taxonomy of CONCOCT bins was then determined by Kraken [29] using the default Minikraken database. BWA 0.7.16 (33) and Samtools 1.3 [34] were used to map the reads back to isolated genomes while CheckM [35] was used to get metrics on the quality of the isolation method.

### Genome annotation and general features

We used QUAST [36] to determine the statistics of the assembly of the *M.* cf. *aeruginosa* CV01 genome. Annotation of the genome was done with PROKKA v1.11 [37] and RAST v2.0 [38] using default parameters to identify putative genes (coding and non-coding sequences). Protein function prediction and annotation of the predicted genes was done against KEGG orthologs (KOs [39] and clusters of orthologs of proteins with Blast2GO [40] and eggNOG v4.5 [41]. Identification of CRISPR repeats, typical in *Cyanobacteria* and in the *Microcystis* genus, was performed with the web server CRISPRFinder [42] and Recognition Tool CRT v1.1 [43], considering a minimum of 3 repeat units. The prediction of transmembrane topology and signal peptide sites was done using Phobius [44] and SignalP v4.1 [45]. Prediction of transmembrane helices in proteins was accomplished using TMHMM v2.0 [46], prediction of ribosomal RNA subunits was done with RNAmmer v1.2 [47], while tRNA and tmRNA genes prediction was done using ARAGORN v1.2.36 [48]. To evaluate the presence and possible origin of prophage sequences, identification and annotation of these sequences were performed using PHASTER [49].

### Detection of toxin genes in *M.* cf. *aeruginosa* CV01

To identify the presence of toxin genes in *M.* cf. *aeruginosa* CV01, we searched the genome to identify protein domains common in toxin gene clusters with Pfam v29.0 [50]. Next, we compared these sequences in our genome with putative toxin synthase genes like microcystin, nodularin, cylindrospermopsin, anatoxin-a, saxitoxin, microviridin, aeruginosin and micropeptin, which are known to be present in cyanobacteria. We used the online tool antiSMASH v3.0 [51] to detect the presence of non-ribosomal peptide synthase (NRPS) and/or polyketide synthase (PKS) gene clusters and other domains of natural products typical of cyanobacterial metabolites. Next, by manually curating the sequences that we identified as being part of toxin synthesis process, we reconstructed the gene clusters ensuring the overlap of contig ends.

### *M.* cf. *aeruginosa* CV01 phylogenetic analysis

To reconstruct the phylogenetic relationships of *Microcystis* strains we inferred a species tree using a set of 12 publicly available fully sequenced genomes. The species tree was constructed from a concatenation of DNA sequences from a set of seven single copy housekeeping genes present in all genomes (*ftsZ*, *glnA*, *gltX*, *gyrB*, *pgi*, *recA* and *tpi*), using maximum likelihood phylogenies to infer the genetic variation of the 12 genomes. Gene sequences were separately aligned with MAFFT v.7.220 [52] and trimmed using Gblocks v0.91b [53,54]. To infer ML phylogenies we used RaxML v8.1.20 [55], to compute and support the trees, by calculating 1000 non-parametric bootstraps using GAMMA distribution and GTR+I+G model, model that was previously estimated using Prottest [56]. Bayesian phylogenies were inferred with MrBayes v3.2 [57] for 1 million generations using the same model and a discarded burn-in rate of 25% of the initial generations. To investigate the relationship between *M.* cf. *aeruginosa* CV01 and other continental African, tropical, and temperate climate strains, we used the phycocyanin alpha subunit and phycocyanin beta subunit (cpcA-cpcB) intergenic space region of the phycocyanin gene cluster (PC-IGS) for a set of publicly available nucleic sequences from *Microcystis* strains with worldwide distribution. Trees were computed using RaxML (1,000 non-parametric bootstraps). Bayesian phylogenies were inferred with MrBayes for 3 million generations. Phylogenies were inferred under a GAMMA distribution and GTR model.

## Results

### Diversity of the microbial communities in the reservoirs

We profiled the bacterial biodiversity in three reservoirs of the island of Santiago using 16S rRNA gene amplicon next generation sequencing. This marker gene is universal in bacteria and it is also present in chloroplasts of eukaryotes, which allowed us to detect both bacteria and eukaryotic phytoplankton. For Poilão reservoir we obtained 804,043 reads (two replicates), for Saquinho 1,271,923 reads (three replicates and Faveta 1,354,273 reads (three replicates), which resulted in 315,291, 437,486 and 425,806 16S rRNA reads for Poilão, Saquinho, and Faveta, respectively.

Regarding taxonomic distribution, Poilão had the highest number of operational taxonomic units (OTUs) with 1,356 OTUs, followed by Saquinho with 966 and Faveta with 950 different OTUs. The OTUs identified in Poilão reservoir represented 39 phyla and included 1.5 % of unassigned sequences. *Proteobacteria* was the numerically most abundant phylum with 76% of all OTUs detected, of which 69.7% were *Gammaproteobacteria,* dominated by the genus *Acinetobacter* (51.6%). The second most abundant phylum was *Bacteroidetes,* representing 4.4% of the OTUs, *Betaproteobacteria* accounted for 3.0% and *Actinobacteria* 2.3%. Cyanobacteria in Poilãao, which represented 3.5%, are mainly represented by the *Phormidium* genus accounting for 2.5% of the OTUs. In Poilão reservoir, sequences assigned to green algae (derived from chloroplasts) were 7.5% of the OTUs and approximately equal amounts of *Chlorophyta* and stramenopiles. Main results regarding the composition of microbial communities are shown in Fig 2.

**Fig 2.**
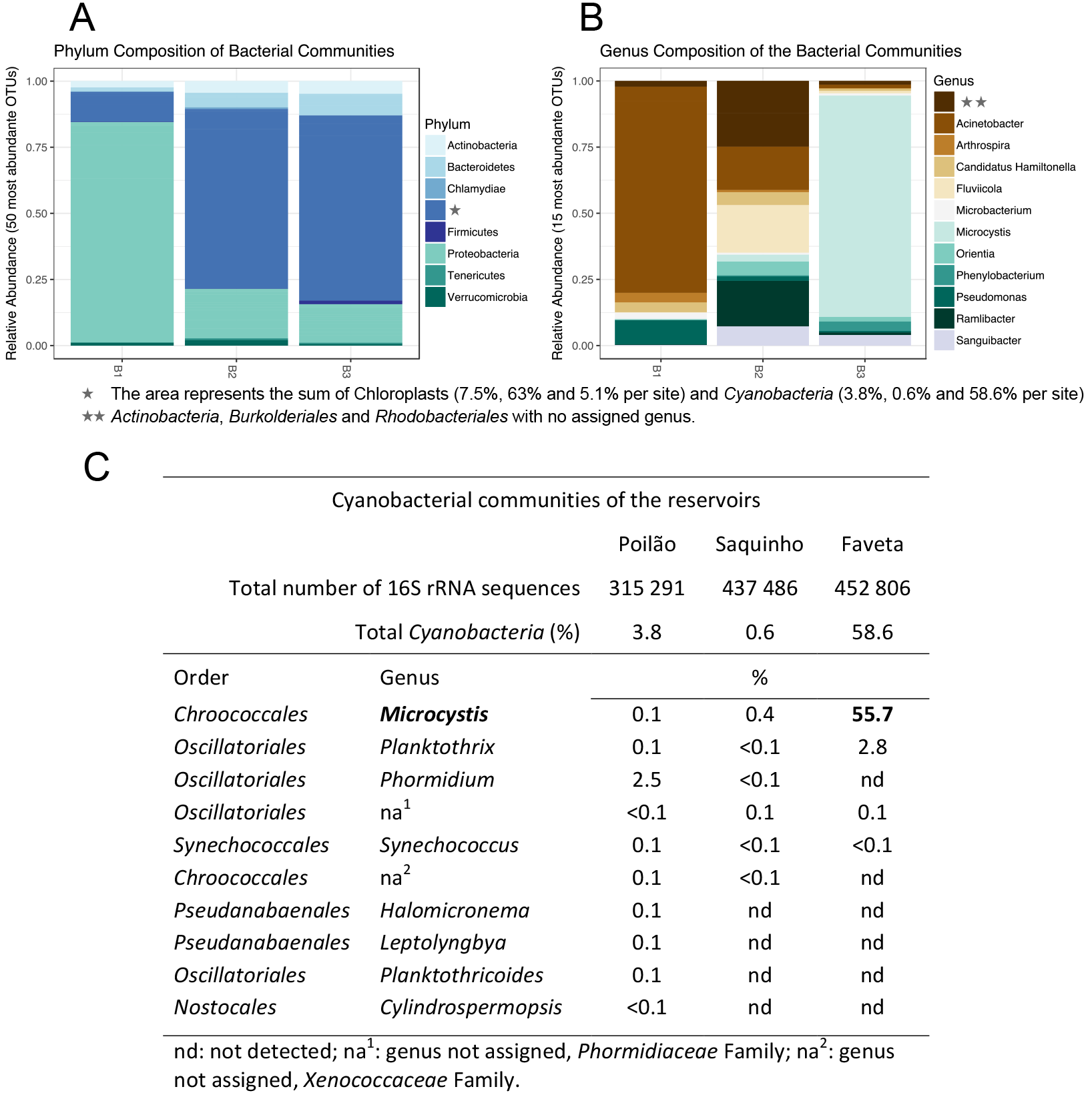
Composition of the microbe communities surveyed in Cape Verde. (A-B)Composition of the microbial communities detected in the three sampled Cape Verdean freshwater reservoirs: Poilãao-B1, Saquinho-B2 and Faveta-B3. (C)Cyanobacteria communities detected in the surveyed reservoirs.

In the Saquinho reservoir, 37 phyla were detected and 0.5% of unassigned sequences. The *Proteobacteria* represented 20%of the OTUs, *Bacteroidetes* 5.4%, *Actinobacteria* accounted for 4.1%, and *Verrucomicrobia* 2.4%. *Cyanobacteria* in this reservoir represented only 0.6% of the OTUs, the lowest percentage of the three sampled reservoirs. This site had the highest number of OTUs associated with green algae (63%), distributed between the *Cryptophyta* with 62% and stramenopiles with 1% of the identified OTUs (Fig 2A). Blastn of the cryptophyte 16S rRNA sequence revealed a hit for *Cryptomonas curvata* plastid partial ribosomal RNA operon (strain CCAC 0006) with 97% identity.

Faveta reservoir had the lowest number of phyla, accounting for 29 distinct phyla and 1.1% of the sequences classified as unknown. The most abundant phylum was *Cyanobacteria* with 58.6% of the OTUs and included two genera known to harbor toxic species: *Microcystis* accounted for 55.7% and *Planktothrix* 2.8% (Fig 2B-C). *Proteobacteria* was the second most abundant phylum with 18.6% assignments; *Bacteroidetes* was next with 7.0% and *Actinobacteria* third with 5.8%: The green algae at this site represented 5.1% of the identified OTUs.

In summary, our survey revealed distinct microbial community structures of the three sampled reservoirs, which is confirmed by the beta diversity analysis in Fig 3D.

**Fig 3.**
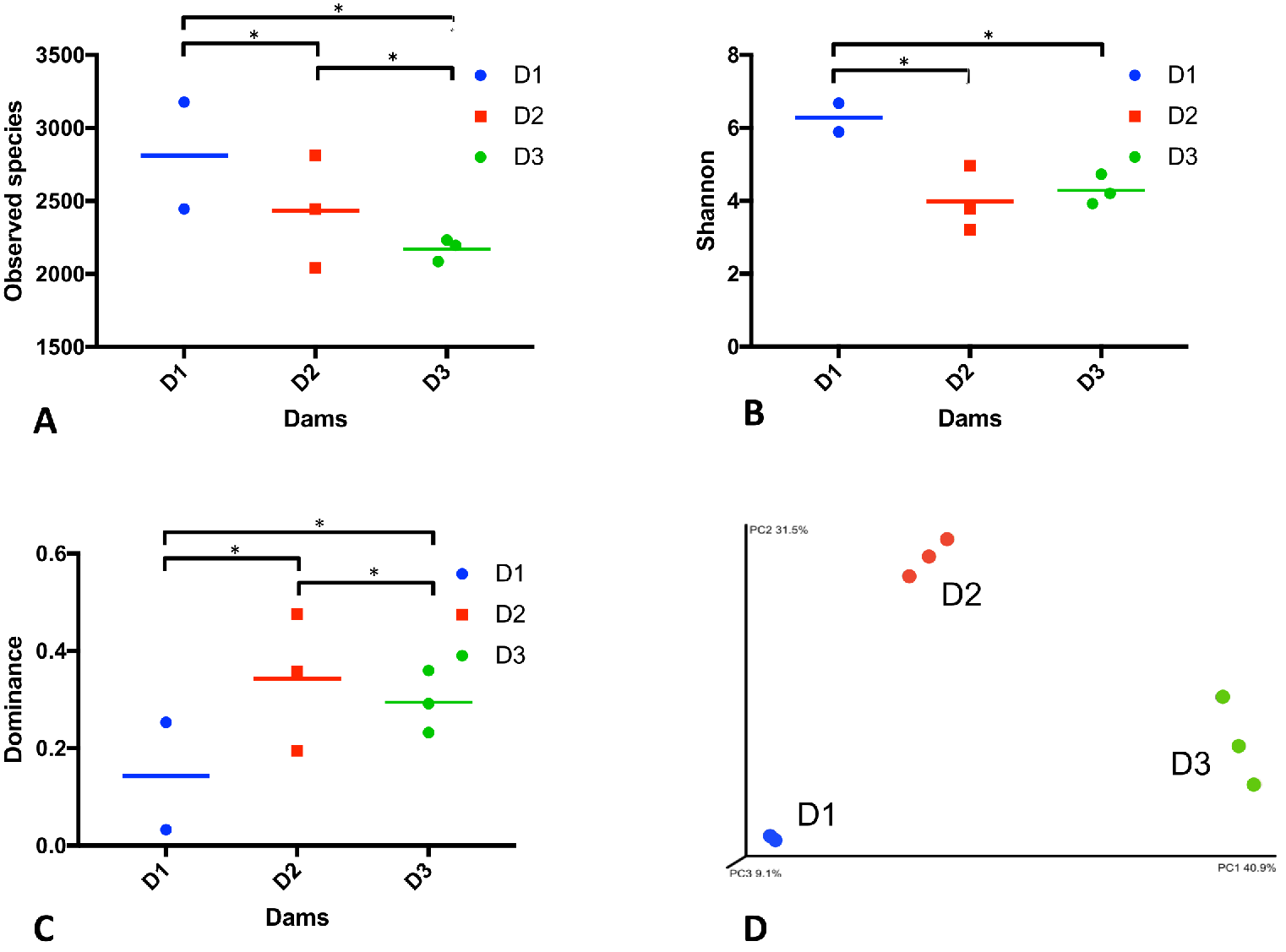
Biodiversity indices per replicate and average. (A) Number of total observed species, (B) Shannon, (C) Dominance and (D) Principal Coordinates Analysis of Beta diversity by weighted Unifrac. Levels of significance: *p < 0.05. Poilão (D1); Saquinho (D2); Faveta (D3).

### Toxin producing genera/species

We identified several cyanobacterial genera in the three sampled reservoirs, some of which are known from the literature to produce toxins. In Faveta reservoir we identified the presence of cyanobacteria from the *Microcystis* and *Planktothrix* genera, reported to produce cyanotoxins, microcystin and anatoxin-a, as well as other potentially toxic molecules like chlorinated and sulfated variants of aeruginosin, cyanopetolin, microginin, and microviridin [58–60]. Poilão reservoir had smaller relative quantities of cyanobacteria, but more genera were identified (Fig 2 and S2 Fig) some of which were potentially toxic like, *Microcystis, Phormidium, Planktothrix* and *Cylindrospermopsis.* These cyanobacteria are known to produce microcystin, homo and anatoxin-a, saxitoxin, and cylindrospermopsin, as well as other metabolites with different levels of toxicity. In Saquinho reservoir the relative quantity of cyanobacteria was the lowest (0.6%, but even so the most represented genus was the potentially toxic *Microcystis*.

### *M.* cf. *aeruginosa* CV01 genome

CONCOCT produced 23 bins, including a bin containing an assembly of *M. aeruginosa* with a 4.7 Mbp genome. According to CheckM with a completeness level of 98.79% with a total of 75.6% of the reads mapping back to its contigs. The SPADES assembled genome was 4.9 Mbp with a completeness level of 99.8% with 84% of the reads mapping back to contig reads from the sequencing effort (S3 Fig and S1 Table). The SPADES assembly was chosen due to its estimated completeness level and total genome size (plotted in a graphical representation in Fig 4), distributed over 262 contigs with length over 1,000 bp and an average GC content of 42.3%. From a total of 5,484 genes annotated, we identified 45 RNA genes, of which 42 were tRNA genes, one tmRNA, and one rRNA set (5S/16S/23S), as well as 5051 protein-coding genes and predicted functions for 91% of all genes (Blast2GO). Considering the importance of metabolite transport and transduction of signals through membranes in cyanobacteria, we also searched for and identified 112 genes with signal peptides and 3,443 genes with transmembrane domains. Major statistical attributes of *M.* cf. *aeruginosa* CV01 genome are described in Table 2.

**Fig 4.**
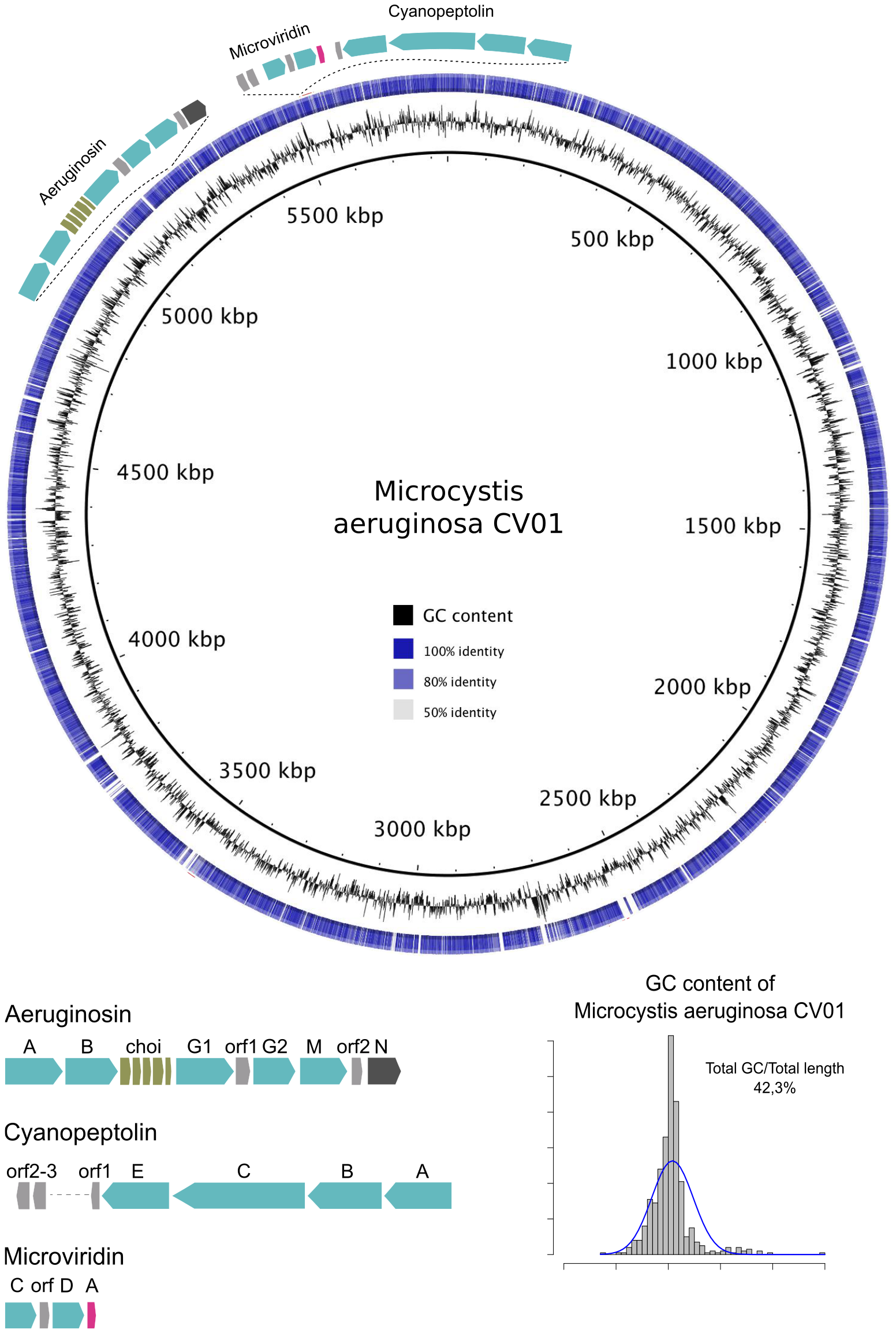
Representation of the genome of *M.* cf. *aeruginosa* CV01, sequenced from the environmental sample of Faveta reservoir. (A) The shades of blue in the circle are indicative of pairwise genomic sequence similarity according to blastn, using as reference the genome of *M. aeruginosa* NIES-843. The location and structure of three gene clusters of potentially toxin-producing secondary metabolites are represented. (B) The histogram represents the GC content of the assembled *M*. cf. *aeruginosa* CV01 genome.

**Table 2.** Genome statistics for *M*. cf. *aeruginosa* CV01.

| Attribute | Value | % of total |
| --- | --- | --- |
| Genome size (bp) | 4,918,369 | 100.0 |
| DNA coding (bp) | 3,900,077 | 79.3 |
| DNA G+C (bp) | 2,080,470 | 42.3 |
| DNA scaffolds | 262 | - |
| Total genes | 5,484 | 100.0 |
| Protein coding genes | 5,051 | 92.1 |
| RNA genes | 45 | 0.8 |
| Genes with function prediction | 4,996 | 91.1 |
| Genes assigned to COGs | 4,166 | 76.0 |
| Genes with Pfam domains | 3,407 | 62.1 |
| Genes with signal peptides | 112 | 2.0 |
| Genes with transmembrane helices | 3,443 | 62.8 |
| CRISPR repeats | 9 | - |

Four partial prophage regions were detected in *M.* cf. *aeruginosa* CV01. The first region, with GC content 45.3%, is 9.4 Kb, encodes for 8 proteins and harbors protein coding sequences (CDS) from a previously described infecting phage, P-TIM68, usually associated with *Prochlorococcus* Myoviridae virus that contains photosystem I gene sequences [61] and transposase sequences. A second partial phage region with 12 CDS and GC content of 43.0% spans over 11.8 Kb and contains phage sequences from the *Microcystis* infecting Myoviridae phage MaMv-DC [62]. Sequences of this phage and of phage Ma-LMM01 [63] are present in a third region with 7.1 Kb in length, 40.4% GC content and 8 CDS. Finally, the forth prophage region of 5.8 Kb in length, coding for 9 proteins and GC content of 40.6% was detected, and CDS from a ssDNA marine virus reported to infect *Synechococcus* [64] were identified. Regarding CRISPR arrays, the defense mechanism of *Cyanobacteria* [65,66] we detected eight CRISPR repetitive units, varying from 0.3Kb and 11.8Kb in length, with direct repeat lengths from 35 to 38 bp. The CRISPR regions and the prophage were not overlapping but two clusters of CRISPR direct repeats (DR were identical to the *Microcystis* phage Ma-LMM01 portion, a memory mechanism to their introduction into the CRISPR locus, providing immunity to further infection by that phage. Other characteristics of the *M.* cf. *aeruginosa* CV01 genome, such as KEGG orthologs and COG functional categories, are summarized in S2-S3 Tables.

### Phylogenetic analysis

Our phylogenetic analysis using seven cyanobacterial genes, as described previously in Materials and methods, identified the dominant cyanobacterium in Faveta reservoir as a *M. aeruginosa* strain, which we named *M.* cf. *aeruginosa* CV01 (Fig 5A). *M.* cf. *aeruginosa* CV01 is placed close to two strains collected in African water bodies: *M. aeruginosa* PCC 9443 collected from a fishpond in Landjica, Central African Republic and *M. aeruginosa* PCC 9807 collected in Hartbeespoort Dam in Pretoria, South Africa (Pasteur Culture Collection of Cyanobacteria. Phylogenetic studies using the PC-IGS intergenic spacer region confirmed the clustering of *M.* cf. *aeruginosa* CV01 with other continental African strains, as shown in Fig 5B, in a branch that contains both microcystin producers and non-producers collected from Ugandan and Kenyan water bodies [67].

**Fig 5.**
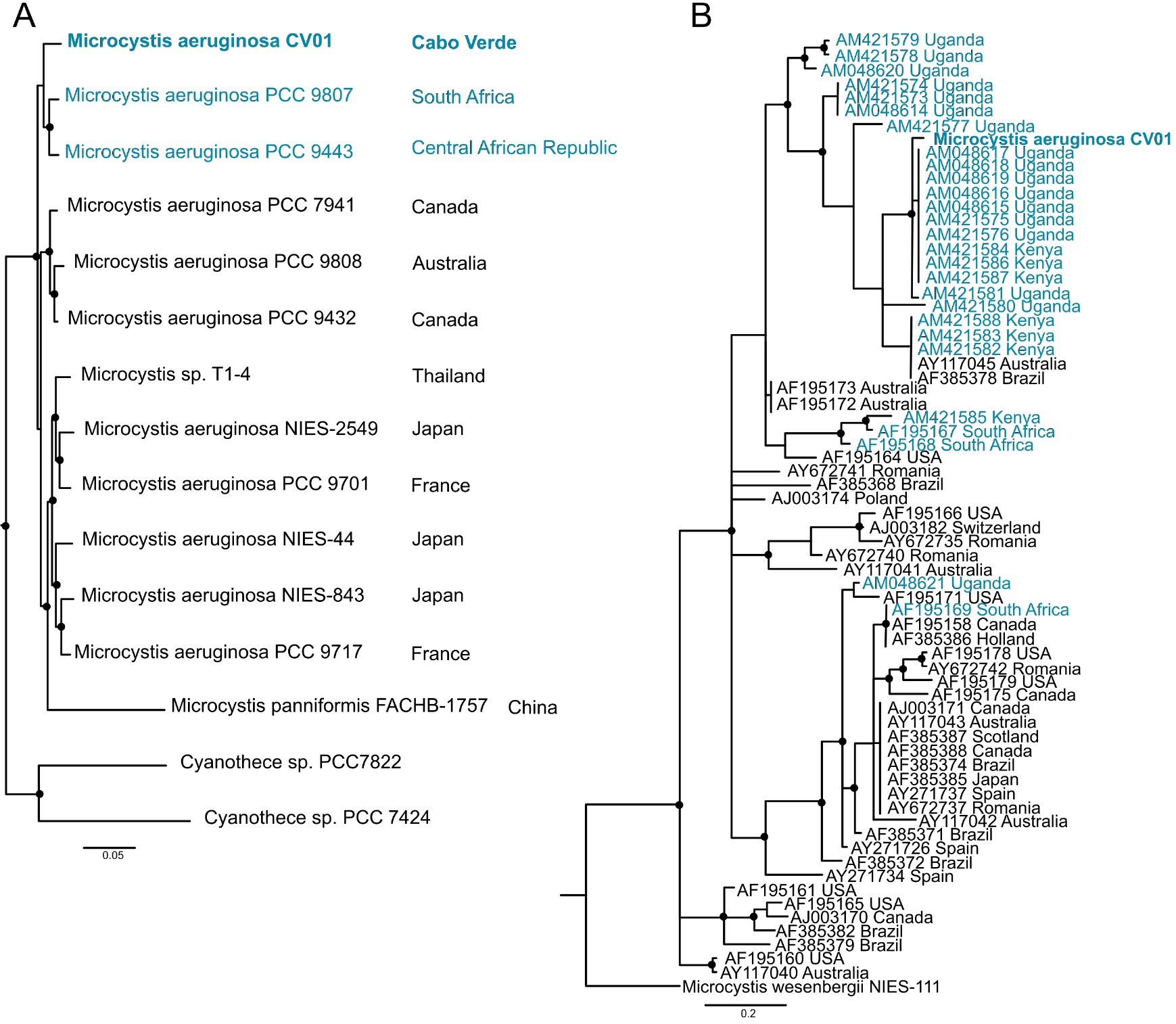
Phylogeny of the Cape Verdean strain *M.* cf. *aeruginosa* CV01. (A) represents the location of *M.* cf. *aeruginosa* CV01 sequenced in this study among a set of fully sequenced genomes of *Microcystis.* The phylogenetic tree was inferred with MrBayes for 1 million generations, using a GAMMA distribution and GTR substitution matrix for a set of 7 housekeeping genes (*ftsZ*, *glnA*, *gltX*, *gyrB*, *pgi*, *recA* and *tpi*). Dots represent posterior probability values higher than 85. (B) A phylogenetic tree of 69 *Microcystis* strains from all over the world locating the Cape Verdean strain among other continental African *M. aeruginosa* strains. ML phylogenies were inferred with RaxML with 1,000 bootstraps and Bayesian phylogenies were inferred with MrBayes for 3 million generations, using a GAMMA distribution and GTR substitution matrix for the intergenic spacer region of the phycocyanin cluster PC-IGS region of 69 globally distributed *Microcystis* strains. Dots represent posterior probability values higher than 75. The African strains are represented in blue in (A) and (B).

### Toxin genes and toxic species

In order to identify the risk of production of toxic cyanobacterial metabolites, we searched the *M.* cf. *aeruginosa* CV01 genome for non-ribosomal peptide synthases (NRPS) and NRPS/polyketyde synthases (PKS) hybrid gene clusters.

Genes that contribute to the synthesis of aeruginosin (NRPS/PKS/saccharide), cyanopeptolin (micropeptin) and microviridin molecules were detected. Aeruginosin and cyanopeptolin gene clusters were present and were non-halogenated (*aerJ* and *mcnD* genes where not detected) (Fig 4). Regarding microcystin gene clusters, some genes were present, nevertheless the full gene cluster was not represented.

Putative genes for the synthesis of cyanopetide metabolites like spumigin (aeruginosin), aeruginoside (aeruginosin), ambiguine (terpene-alkaloid), and piricyclamide (post-translational modified peptides) were also detected, but not their complete gene clusters. Finally, one unknown NRPS/PKS gene cluster was identified, showing the potential to produce peptide molecules that are yet unknown.

### Nucleotide sequence accession numbers

The nucleotide sequence data are available at DDBJ/EMBL/GenBank under the accession number SUB2733390.

## Discussion

The studied reservoirs show three distinct microbial/microalgae community profiles, despite being located on the same island and in a radius of 15 kilometers from each other: in two sites, bacteria were dominant (*Proteobacteria* and *Cyanobacteria*) and in the other reservoir microalgae belonging to the cryptophytes were the most abundant taxa. The dominant species from one of the reservoirs was identified as a *M. aeruginosa* strain through phylogenetic studies, placing it closer to other strains collected in continental Africa. *Microcystis* spp. were detected in all three reservoirs, as well as other cyanobacteria known to bloom and produce cyanotoxins. Our analysis of the assembled *M.* cf. *aeruginosa* CV01 genome revealed that it can produce toxins and therefore, a potential risk of toxin production can exist in Cape Verdean reservoirs.

Our microbial biodiversity survey revealed distinct community structures in the three reservoirs (Fig 3D). In fact, *Proteobacteria* dominated Poilão reservoir, mainly bacteria from the *Acinetobacter* genus, while in Saquinho reservoir microalgae from the *Cryptomonas* genus were dominant, and in Faveta reservoir the cyanobacterial strain *M.* cf. *aeruginosa* CV01 was the most abundant. Despite being located on the same island and having common microbial groups, we found that each phyla’s quantitative distribution varied substantially between reservoirs (Fig 2A). Nevertheless, the profiles from the reservoirs identify groups that are common in other water bodies studied worldwide despite the differences in relative quantities as is the case of *Cyanobacteria*, *Cryptophyta*, *Actinobacteria*, *Bacteroidetes*, *Verrucomicrobia*, and the three clades of *Proteobacteria* (Alpha, Beta and Gamma) [14,68]. Actually the microbial profile of Poilão reservoir resembles those of lakes close to urban areas, where *Acinetobacter* is the dominant genus [69].

In each reservoir, one genus prevailed with a relative abundance above 50%: in Poilãao was this was *Acinetobacter,* in Saquinho it was *Cryptomonas,* and in Faveta the *Microcystis* genus.

The analysis of the local diversity indices of the replicates from each reservoir revealed consistency and reinforced the sites’ distinct microbial communities (Fig 3A-C). The indices also showed that microbial communities presented different dominant genera in each of the reservoirs, as well as abundance of different taxa in all sites as typically found in freshwater bodies around the world [7–10,12].

Besides the operational starting date differences between the reservoirs (Table 1) and no physical communication between lakes, abiotic factors specific for each site might be involved in the dominance variations within the microbe communities.

The dominance of one cyanobacterial strain in Faveta, allowed us to assemble and fully study its genome, and to identify genes, allowing the reconstruction of toxin pathways and assessing the toxin risk inherent in this specific strain. Exploration of the assembled genome also revealed genomic features in common with other *M. aeruginosa* genomes (Table 2). Some phage genes were found integrated in the genome of *M. cf. aeruginosa.* Myoviridae “photosynthetic” freshwater cyanophages (Ma-LMM01 and MaMV-DC) were also found. These genes are thought to play an important role during phage infection by supplementing the host with the production of photosynthesis proteins, a process that can be also beneficial to the host during the infection process as suggested by some authors [61,70,71]. These horizontal gene transfer events are shaping the genome architecture of the *Microcystis* genus, providing a supplementary advantage that can be important during cyanobacteria blooms. A region containing chlorophyll a apoproteins A1 and A2 synthesis genes was also identified, but since these are single copy genes located near transposase sequences in this new genome, they were probably misidentified as having phage origin.

We identified four NRPS/PKS gene clusters that could synthesize potentially toxic metabolites: three well-known metabolites (aeruginosin, cyanopeptolin, and microviridin, represented in Fig 4A) and another metabolite from one yet unknown gene cluster. The real toxic potential of these metabolites is difficult to determine, as the actual synthesis pathways are not fully known, so further toxicological screenings need to be performed.

Phylogenetic markers placed the Cape Verdean strain among others from freshwater bodies from Africa, albeit Cape Verde being a distant archipelago from the continental Africa. The identification in all reservoirs of other cyanobacterial genera known to be toxin producers like *Phormidium*, *Planktothrix*, *and Cylindrospermopsis*, increases the potential risk of toxin production. Other studies in African water bodies have identified these and other potentially cyanotoxin producers, raising the possibility of future occurrence of other cyanobacterial genera in Cape Verdean freshwater reservoirs.

Cyanobacterial blooms occur in freshwater reservoirs distributed worldwide where *M. aeruginosa* is one of the most frequently detected species. Actually, many long-term studies have reported toxic blooms in lakes and rivers from Kenya, Uganda, Senegal, Morocco, and South Africa [67,72–77], often dominated by *M. aeruginosa,* as we also detected in the island of Santiago. Moreover a recent review on the occurrence of cyanobacterial blooms in Africa [78] shows that there is limited information from western African countries, including Cape Verde, exposing the need for further studies in countries were water quality is threatened and scarce. Therefore, our study increases the available information on cyanobacterial communities described for the western African region. The scarcity of renewable freshwater resources of archipelagic states like Cape Verde is aggravated by the terrain that favors torrential water flows and strong anthropogenic pressures on the environment leading to eutrophication of its freshwater bodies and increased risk of toxic algal blooms.

The main threat and concern from our analysis, was the identification of a bacterial community dominated by *M*. cf. *aeruginosa* CV01, which signals the possibility of toxic blooms in Cape Verdean reservoirs, since exponential growth is typical of this species.

The occurrence of blooms and toxin production are potential life-threatening risks to public health, so monitoring plans are very important. The costs involved in these control and containment strategies can be prohibitive especially for low and middle-income countries. Hence, approaches like the one proposed in this work, which enabled the identification of potentially toxic cyanobacterial genera through 16S rRNA gene markers could be an interesting alternative, without the time-consuming and expertise-dependent microscope identification of toxin-producing organisms or mass spectrometry-base identification of toxins. NGS is still not widely available but DNA sampling kits are easy to use and can be sent to sequencing facilities at cost-effective prices. Other more sophisticated technics are possible such as lab on a chip, mass spectrometry or even portable NGS devices, that can be adapted to use our workflow in the field, but if a simple molecular lab is available, a PCR assay could also be efficient to detect the presence of specific putative cyanotoxin genes. These strategies can alert authorities and populations before bloom formation and toxin production.

Cyanobacteria are currently being developed and used for bio-production of metabolites and biomass in algal farms, for example in the production of human dietary supplements, fertilizers, animal feed or biodiesel, to name a few uses [79–83]. Cape Verde has little land suitable for agriculture, but it has temperature and light conditions that are speculated to be suitable for simple bioreactors for biomass production using photosynthetic microorganisms like green algae or cyanobacteria. The observation of spontaneous blooms of cyanobacteria in freshwater reservoirs lends credibility to this hypothesis, and opens the way for a new productive industry in the archipelago.

The present work identifies the existence of real risk for cyanotoxin production in Cape Verdean freshwater reservoirs. Similar structures are planned, which will also need to be studied and monitored. Future work should include studies on the dynamics of the local microbial communities, as well as characterize how environmental factors are affecting their organization, in order to predict and control the impacts of water impairment and toxin production on public health and on the economy. In this study we made use of many freely available open source tools, which represent, to our knowledge, innovative research strategies in Cape Verde. This study will open the way for further research on microbial biodiversity and other genomic studies in the archipelago, and raise questions relevant for different areas of research and application.

## Aknowledgements

We wish to thank Dr. W. Szymaniak for help in accessing the sample sites and the Gene Expression Facility from Instituto Gulbenkian de Ciência for all the technical support. Ana P. Semedo-Aguiar was recipient of fellowship SFRH/BD/113752/2015 from the Portuguese Science and Technology Foundation (FCT), under the auspices of the Programa de Pós-Graduação Ciência para o Desenvolvimento (PGCD) and RBL was recipient of fellowship SFRH/BPD/91518/2012 from FCT.

## Author Contributions

Conceived and designed the experiments: APSA JBPL RBL. Performed the experiments: APSA RBL. Analyzed the data: APSA RBL. Contributed reagents/materials/analysis tools: JBPL RBL. Wrote the paper: APSA JBPL RBL.

## Supporting Information

**S1 Fig. - The GC content of the raw reads and the final assembled metagenome from Faveta reservoir**. GC content of the raw reads and of the assembled genomes in the histograms of panels A and B, respectively.

**S2 Fig. - Rarefaction curves per number of reads for each reservoir sampled**. Representation based in Chao1, Observed OTUs and Phylogenetic distance - PD whole tree of the 16sRNA gene amplicons of Illumina sequencing. Dl-Poilão; D2-Saquinho; D3-Faveta.

**S3 Fig. - Faveta reservoir**. Degree of completeness and contamination of the *M. aeruginosa* bin detected using CONCOCT and assembly produced by SPADES (panel A), Graphical representation of a PCA of clusters based on composition and coverage (bin 19 in yellow). Left Panel C - Histogram of GC distribution of sequences within bin 19, Right Panel C - Deviation of the average of the entire assembly, x-axis represents GC and y-axis the size. Left Panel D - Histogram of Manhattan distance between tetranucleotide signature of bin 19 and the entire assembly, Right Panel D - Deviation of the average of the entire assembly, x axis represents GC and y axis the size.

**S1 Online Fig. - Kraken analyses of taxon distribution in whole genome reads and 16S rRNA from Faveta reservoir.**

## Supplementary tables legends

**S1 Table. Degree of completeness and contamination and percentage of mapped reads**

**S2 Table. KEGG functional annotation of the predicted transcriptome of *Microcystis* cf. *aeruginosa* CV01**

**S3 Table. COG functional categories of the predicted transcriptome of *Microcystis* cf. *aeruginosa* CV01**

